# Single-cell RNA-seq data reveals TNBC tumor heterogeneity through characterizing subclone compositions and proportions

**DOI:** 10.1101/858290

**Authors:** Weida Wang, Jinyuan Xu, Shuyuan Wang, Peng Xia, Li Zhang, Lei Yu, Jie Wu, Qian Song, Bo Zhang, Chaohan Xu, Yun Xiao

## Abstract

Understanding subclonal architecture and their biological functions poses one of the key challenges to deeply portray and investigative the cause of triple-negative breast cancer (TNBC). Here we combine single-cell and bulk sequencing data to analyze tumor heterogeneity through characterizing subclone compositions and proportions. Based on sing-cell RNA-seq data (GSE118389) we identified five distinct cell subpopulations and characterized their biological functions based on their gene markers. According to the results of functional annotation, we found that C1 and C2 are related to immune functions, while C5 is related to programmed cell death. Then based on subclonal basis gene expression matrix, we applied deconvolution algorithm on TCGA tissue RNA-seq data and observed that microenvironment is diverse among TNBC subclones, especially C1 is closely related to T cells. What’s more, we also found that high C5 proportions would led to poor survival outcome, log-rank test *p*-value and HR [95%CI] for five years overall survival in GSE96058 dataset were 0.0158 and 2.557 [1.160-5.636]. Collectively, our analysis reveals both intra-tumor and inter-tumor heterogeneity and their association with subclonal microenvironment in TNBC (subclone compositions and proportions), and uncovers the organic combination of subclones dictating poor outcomes in this disease.

**Highlights:** We applied deconvolution algorithm on subclonal basis gene expression matrix to link single cells and bulk tissue together.

## Introduction

Triple-negative breast cancer (TNBC) is one special type of breast cancer which does not express the genes for estrogen receptor (ER), progesterone receptor (PR) and human epidermal growth factor receptor 2 (Her2), and always accompanied with weaker treatment effect and worse clinical outcome (Lehmann et al., 2011; Li et al., 2015; Liu et al., 2018; Marra et al., 2019). Most triple-negative breast cancers have a basal-like genetic pattern (ER- / PR- / Her2-), and It has been reported that 56–85% of TNBCs are basal-like breast cancers (Lachapelle and Foulkes, 2011; Li et al., 2015; Prat et al., 2013). What’s more, TNBC is a disease with heterogeneity not only inter-tumor but also intra-tumor, and represents various clinically and biologically distinct subgroups that have not been clearly defined yet (Cho et al., 2011; Karaayvaz et al., 2018; Mills et al., 2018; Montagna et al., 2013).

Inter-tumor heterogeneity denotes that patients suffering from one same cancer but have a great difference on clinical outcome and factors that can cause this phenomenon are various, such as gene expression, mutation events, methylation pattern and so on (Cusnir and Cavalcante, 2012; Nault, 2014; Wright et al., 2017). And nowadays, tumor microenvironment has arisen more interests which generally means non-cancerous cells present in and around a tumor (Buoncervello et al., 2019; Ferrone and Whiteside, 2007; Zheng and Gao, 2019), and especially, tumor immune microenvironment which focuses on immune cells has become one research hotspot (Bao et al., 2019; Jiang et al., 2019; Zheng et al., 2019). On the other hand, intra-tumor heterogeneity denotes that some subpopulations of cancer cells which differ in their genetic patterns, phenotypic characteristics or cell type composition within a given tumor tissue (Cresswell et al., 2016; Stanta and Bonin, 2018; Wright et al., 2017). Each subpopulation of cancer cells would build up a distinct subclone and if get a growth advantage, that subclone population can expand over time and inhibit proliferation of normal cells (Giessler et al., 2017; Newcomb et al., 1978). While in this work, we considerate subclone as one special existence that subclonal compositions can reveal intra-tumor heterogeneity and subclonal proportions also can reveal inter-tumor heterogeneity. Hence, we believe that analyze subclonal populations can help us to deeper understand tumor heterogeneity in TNBC.

Meanwhile, deep sequencing techniques are experiencing in high-speed development and the appearance of single-cell sequencing technologies transfers researchers’ perspective from traditional bulk tissue samples to single cells (Aran et al., 2019; Chu et al., 2016; Schelker et al., 2017; Zong, 2017). Armed with that technique, researchers can analyze every single cell on features of somatic mutations, copy number variations, gene expression, methylation and so on, then mine deeper differences among distinguish cell populations (Chen et al., 2016; Garvin et al., 2015; Hui et al., 2018; Poirion et al., 2018). For example, Olivier et al. developed a linear modeling framework, SSrGE, combined with single-cell sequencing to link eeSNVs associated with gene expression, and further identified different subpopulation and linkage of genotype-phenotype relationship (Poirion et al., 2018). Anna et al. developed a novel method: Topographic Single Cell Sequencing (TSCS) to measure genomic copy number profiles of single tumor cells, and applied it to 10 early-stage breast cancer patients. And they found that multiple clones crossed the basement membrane that brings out the invasion in Ductal carcinoma in situ (Casasent et al., 2018).

There are only a small number of studies have characterized the intra-tumor heterogeneity of TNBC through single-cell RNA-seq revealing the distinguish patterns that can reflects evolution of copy number variations during tumor progression. While these findings still focus on signature gene markers of cell subpopulations (Aarts et al., 2001; Arienti et al., 2017; Karaayvaz et al., 2018), but in our study we mainly analyze subclonal proportion in tissue bulks. Based on the deconvolution algorithm (Jin et al., 2018; Newman et al., 2015; Zeng et al., 2019), we connected single-cell level with bulk-tissue level by estimating subclonal proportion in tumor tissue then reveal genetic features, biological functions and clinical behaviors of TNBC subclones. And in result, we found that distinct subclone has its special function such as “regulation of T cell activation” in C1, “regulation of ARF GTPase activity” in C3, “programmed cell death” in C5 and so on.

## Results

### Subclone within malignant TNBC tumor blocks

Tumor subclones usually tend to have different mutation patterns and copy number distributions, resulting in a difficult task in the identification of relative tumor subclone-specific clusters. Hence, we used “inferCNV” methods to estimate single-cell CNV profiles from RNA-seq read counts through comparing with GTEx (https://www.gtexportal.org/home/) normal breast tissues which was used as reference. After that, we applied k-means clustering methods on that inferred CNV data which were log2-transformed and distinct subclones within 1534 single cells. At first, we used “clusGap” function with default default arguments to estimate the number of clusters and we found split to 5 representative clusters was most ideal for our data. Then to determine the relatively optimal seed, we clustered 1000 times (set seed from 1 to 1000) and compared tot.withinss (the total within-cluster sum of squared errors) and betweenness generated from each time. Finally we set 34 as the best seed which has the lowest tot.withinss and largest betweenness. Combined with the above number of clusters and seed, we finally distributed every single cell to its closest group in 5 clusters (Figure 1A). What’s more, we also identified 767 malignant cells in those cells which have accumulated sum of absolute CNV (log2-transformed) over median values. Then to verify the classification result, we compared tumor scores transformed from ESTIMATEScore [(max(x) – x) / (max(x) - min(x))] in those cells with normal cells and found they have significant Wilcoxon Rank Sum *p*-value < 2.2e-16 (Figure 1B, 1C) (Schelker et al., 2017).

**Figure 1.**
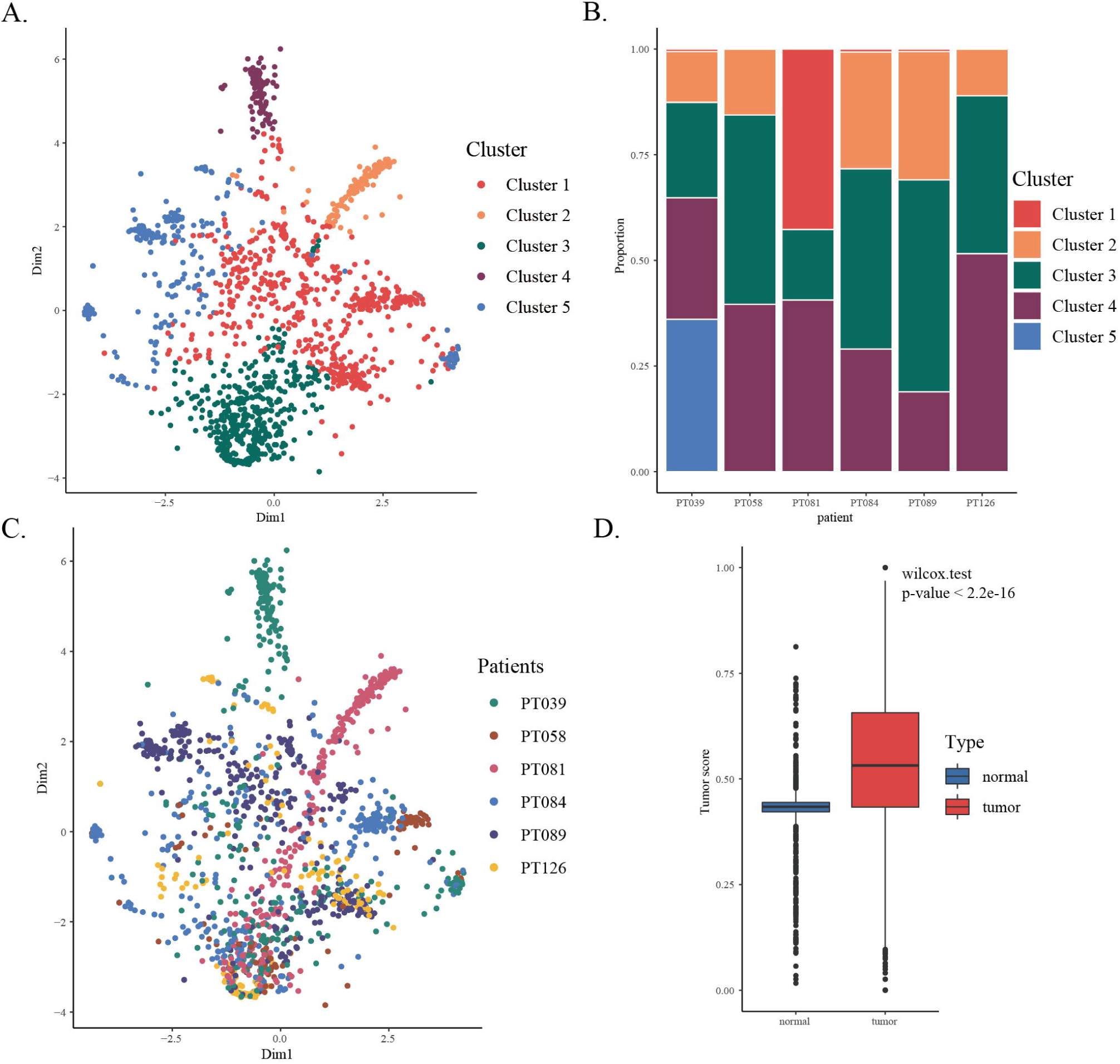
Subclone identification based on single-cell CNV and malignant cells classification. **A** T-SNE plot for 1534 single-cells and their distributions in five subclones. **B** Bar plot depicting the proportions of five subclones in patients. **C** T-SNE plot for 1534 single-cells and their distributions in six patients. **D** Box plots representing the distribution of tumor scores between malignant cells and normal cells.

After that, we identified distinct cell populations within the tumors by comparing with reference HPCA dataset (Human Primary Cells Atlas) as described in Methods. We used 153 primary cell types to identify our single cells, and 29 main cell types were matched. And we found 26.01% single cells and 35.3% malignant cells (Figure 3F) are more likely to be induced pluripotent stem (iPS) cells that maybe indicated most cancer single cells have pluripotent potentiality (Malchenko et al., 2010; Werbowetski-Ogilvie and Bhatia, 2008). Meanwhile, 21.71% cells were similar to neutrophil may hinted that tumor tissue could recruit neutrophil and cause inflammatory response. While except for the above two type, none of the rest cell types’ proportion was more than 10%. On the other hand, for each cell cluster, we also statistics cell types within and found that both C1 and C5 have more than 40% cells were iPS cells, and in C2 and C3 iPS cells’ proportion also the highest (28.40% in C2 and 24.43% in C3), but only C4 have 58.75% cells were Neutrophils (Figure 3).

**Figure 2.**
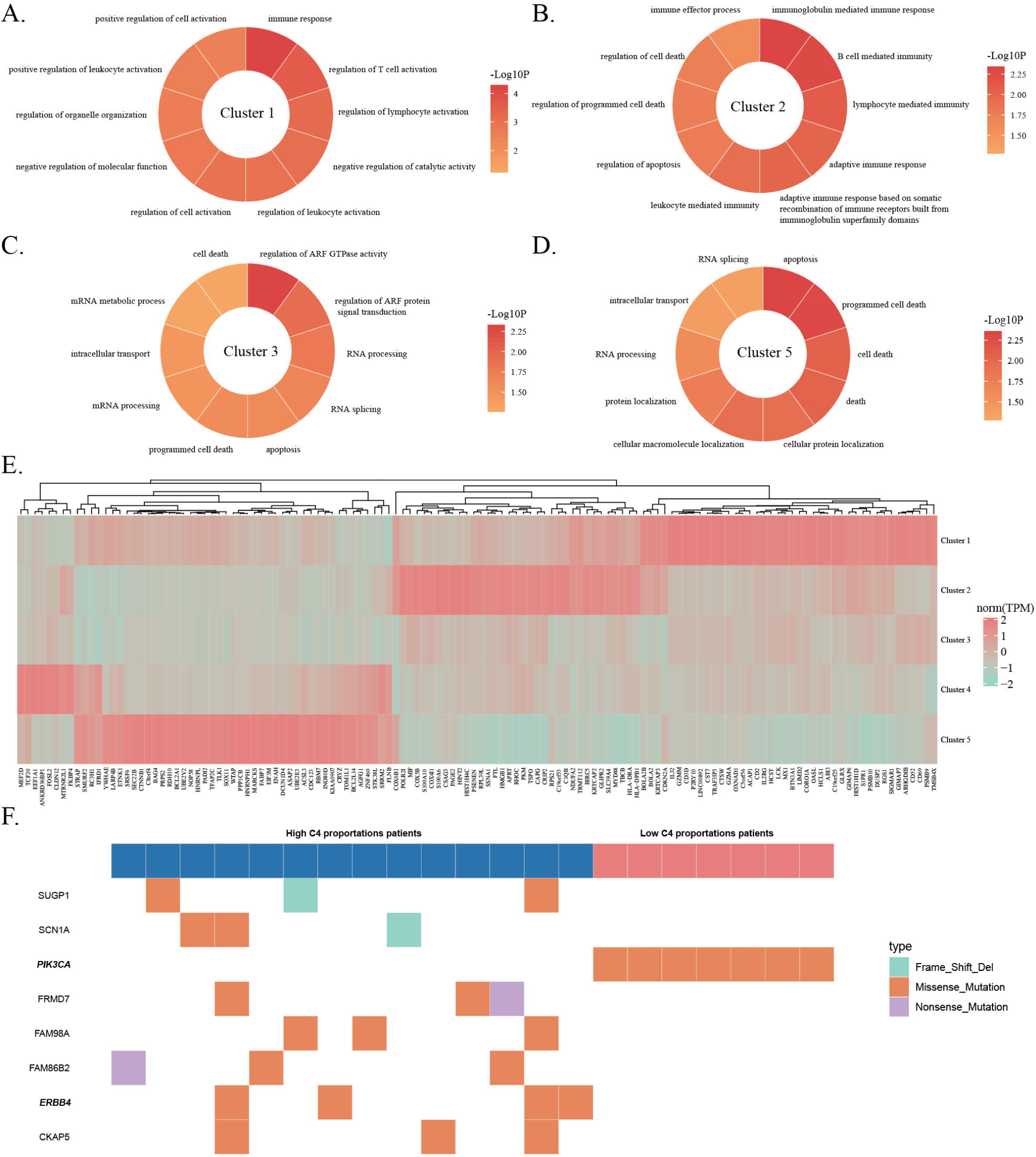
Subclonal GO terms enrichment and heatmap for gene markers expression. **A - D** pie plot depicting the enriched top 10 Go terms in cluster 1, 2, 3 and 5. **E** heatmap for gene markers expression in five subclones. **F** differential mutated genes in cluster 4.

**Figure 3.**
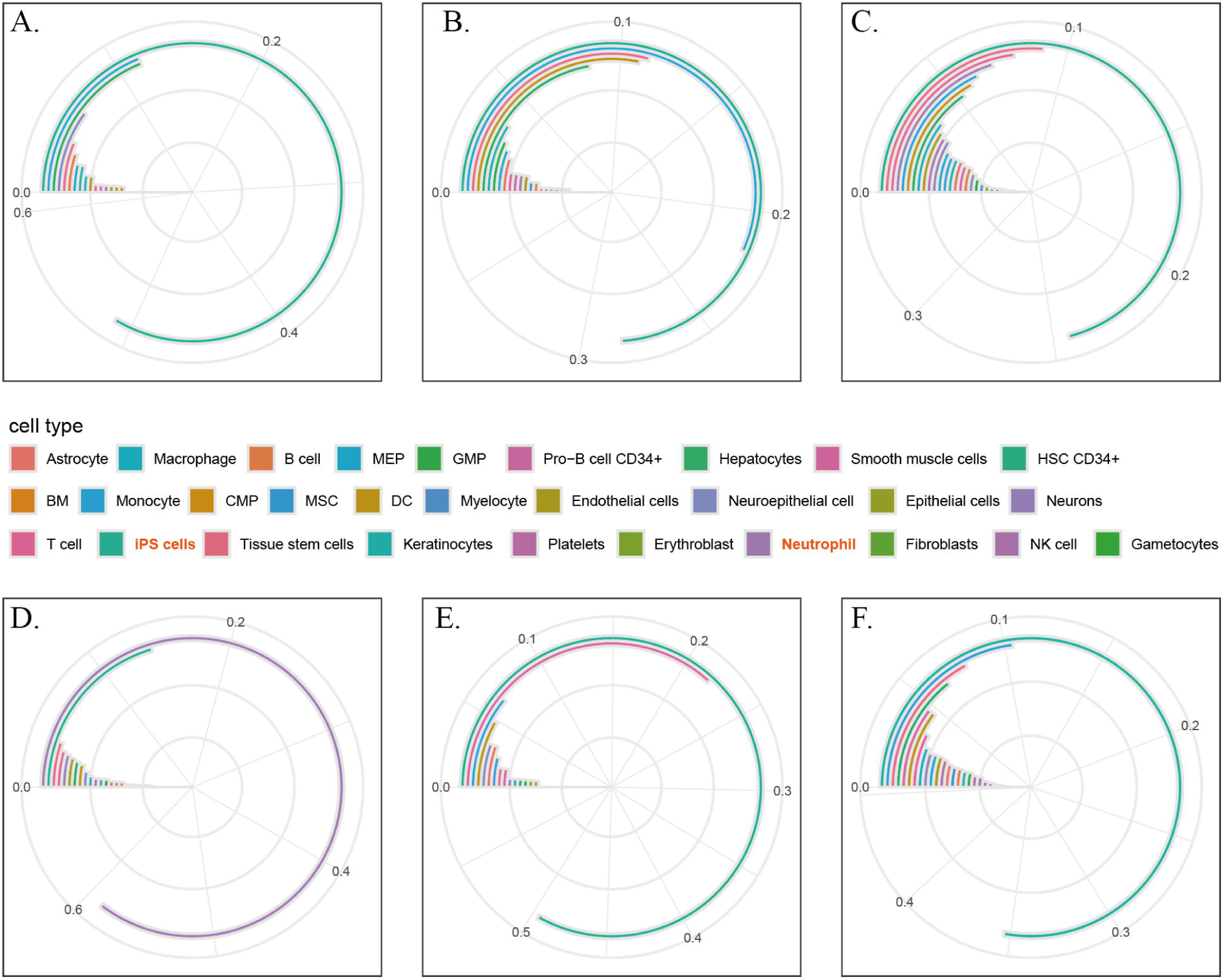
Cell types identification and their distributions in five subclones. **A - E** depicting the cell type proportions in five subclones. **F** representing the proportions in malignant cells.

Considerate on that gene expression profile data is more widely used and easier to get, hence, we characterized the above 5 subclones on gene expression data and construct a basis matrix to deconvolute on TCGA bulk tissues (Figure 2E). From cell to tissue, we transformed TCGA bulk gene expression matrix to subclone proportion matrix, and through this way we reduced the dimension from tens of thousands genes to only just five subclonal proportions. Then based on subclone proportions, we can reveal how those subclones influence tumor development and what’s role they played in by analyzing the changes in subclonal composition among patients.

### Heterogeneous subclones play different roles

With further research, we found that every subclone plays important but entirely different roles, and the organic combination of those subclones makes tumor so complicated and uncontrollable. And we hypothesized that inter-tumor heterogeneity, in some instance, was affected by intra-tumor heterogeneity, especially the compositions of subclones.

#### C1 and C2 associated with immunity

Based on highly expressed gene markers identified above, we annotated GO BP terms for C1 and C2 clusters by DAVID and found that two subclones may be tightly associated with immune-related functions.

For C1 subclone, we got one GO term “regulation of T cell activation” indicated C1 subclone may closely relate to T cells (Figure 2A). To verify the relevance between them, we estimated both subclone and immune microenvironment in TCGA TNBC patients by CBS deconvolution algorithm provided in “EpiDISH” package, and then used Wilcox Rank Sum Test to check the proportion difference of 22 special immune cells (CIBERSORT LM22 dataset) among patients with high and low (threshold was mean value) C1 proportions (Newman et al., 2015). We observed that CD4 T cells, Treg cells, Macrophages and mast cells have significant difference (with Wilcox Rank Sum Test *p*-values = 1.2846e-5, 0.0024, 5.1568e-07 and 0.0036, Figure 4A) which may indicated that C1 subclone may influence immune-related function via changing T cell microenvironment. What’s more, we also checked whether patients with high and low C1 proportion have different immune ability. We used “EASTIMATE” package to calculate immuneScore for each TNBC sample and found that high C1 proportion patients have significant higher scores than low C1 proportion (Wilcox Rank Sum Test, greater alternative, *p*-value = 0.0478, Figure 4B).

**Figure 4.**
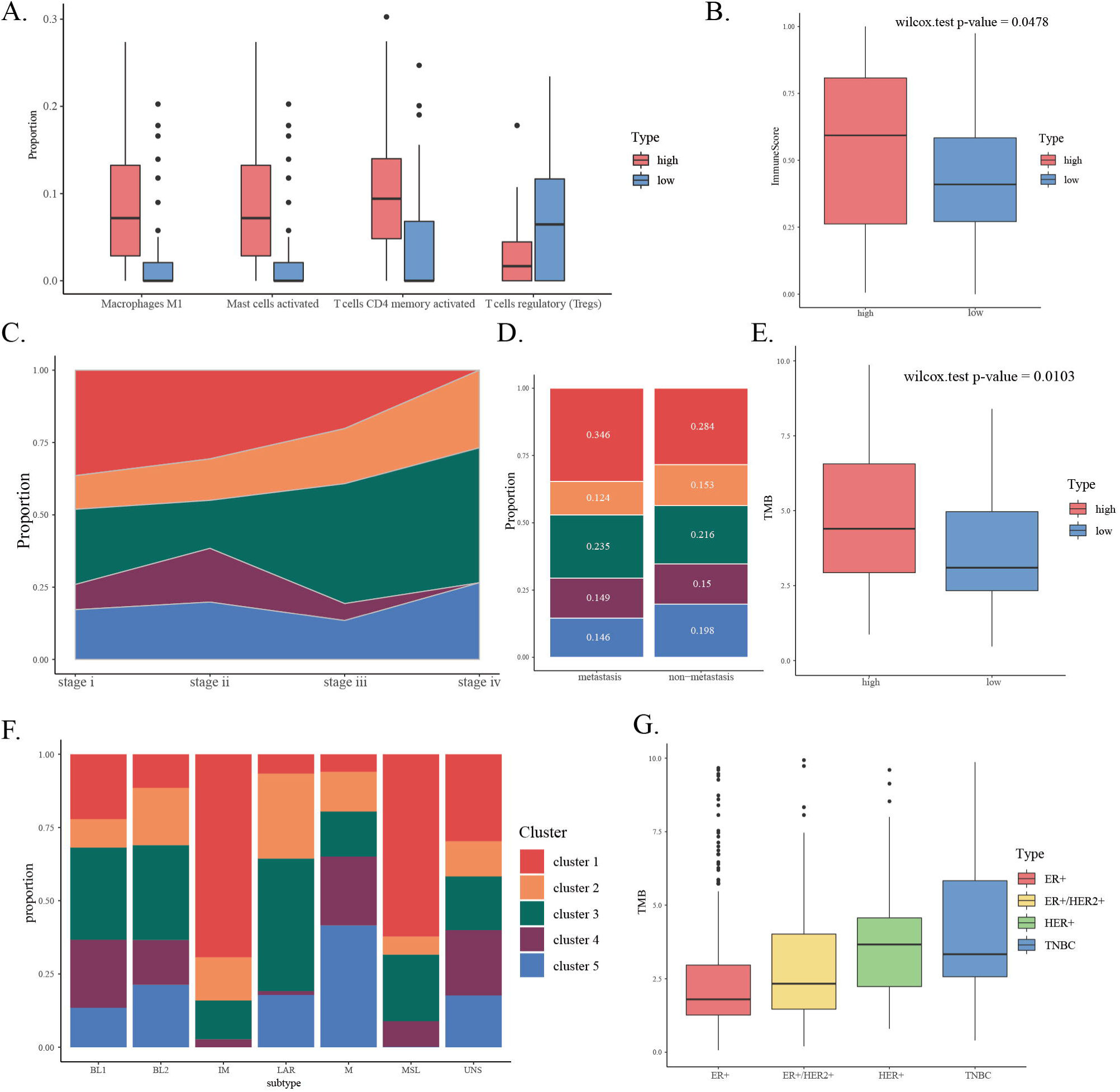
Analyses focus on cluster 1, 2 and 3. **A** depicting differential immune cell types between patients with high and low C1 proportion. **B** boxplot for ImmuneScore between patients with high and low C1 proportion. **C** representing the distribution of five subclones in different stage patients. **D** representing the distribution of five subclones in metastatic and non- Metastatic patients. **E** boxplot for TMB between patients with high and low C2 proportion. **F** representing the distribution of five subclones in TNBCtype6. **G** boxplot for TMB among ER+, ER+/HER2+, HER+ and TNBC patients.

As for C2 subclone which was related to GO term “lymphocyte mediated immunity” and “immunoglobulin mediated immune response” (Figure 2B), while we observed that there was no significant difference between high and low proportion patients in immune microenvironment and immune ability. But very interestingly, we found that high C2 proportion patients have higher Tumor Mutational Burden (TMB) than low C2 proportion (Fold Change was 2.110048 and Wilcox Rank Sum Test *p*-value was 0.01027, Figure 4E), showing that high C2 proportion patients may have benefits from immuno-oncology (IO) therapies (Allgauer et al., 2018; Boumber, 2018; Romero, 2019; Wu et al., 2009).

Furthermore, we also compared TMB among four main IHC breast cancer subtypes (including ER+, HER2+, ER+ / HER2+ and TNBC), and we found that both HER2+ and TNBC have more higher TMB than the other two, and TNBC compared with ER+ and ER+ / HER2+ were all significant (Liu et al., 2018; Marra et al., 2019) (Wilcox Rank Sum Test *p*-value were 7.841e-13 and 2.525e-4, Figure 4G).

#### C3 may bring about metastasis

Different from others, C3 subclone has none significant highly expressed genes, so we used significant down-expressed genes to characterize its biological functions and roles in tumor development.

We used DAVID to annotated GO BP terms, and found that C3 subclone related to “regulation of ARF GTPase activity” and “regulation of ARF protein signal transduction” (Figure 2C). While some researchers proposed that ARF GTPase-activating proteins may play an important role in tumor metastasis (Campa and Randazzo, 2008; Hongu et al., 2016). On the other hand, we also found that C3 proportion abnormal rise in stage III and IV patients, maybe it plays an important role in conversion from stage II to III and IV, especially maybe associated with tumor metastasis (Figure 4C). To support that, we collected micrometastasis information and first non-lymph node metastasis anatomic sites of TNBC patients, and tested whether high C3 proportion interrelated with tumor metastasis.

We used one-side Wilcoxon Rank Sum Test to check whether patients with high C3 proportion really have higher probability to suffer from tumor metastasis, but we got *p*-value was 0.542. Hence, we thought that there is not one subclone plays a role in tumor metastasis process but subclonal cooperation drives metastasis. Then we compared the subclonal distributions between metastatic patients and non-metastatic patients, we found that, in metastatic patients, C1 and C3 raise, C2 and C5 declined, only C4 were stable (Figure 4D). Hence we hypothesis that C1 / C3 were risk factors and C2 / C5 were protection factor, then we constructed a risk score to divide samples into high or low risk of metastasis (threshold is MAD), and we verified low risk score generally related to low risk of metastasis (Cumulative Hypergeometric Test *p*-value was 0.0594, Table 2).

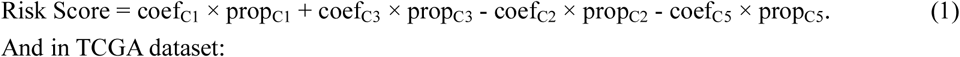

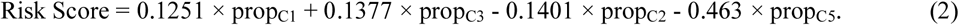

Where coef were coefficients got from cox proportional hazards regression model of subclonal proportions for 5-years TNBC patients, and prop were the proportions of subclones.

#### C4 may be constructed by neutrophils

When we annotated functions for differential expression genes in C4, we found that there is no GO term or KEGG pathway enriched in neither up- nor down-regulated gene sets. But we found that a great part of cells in C4 (44%) have a similar expression pattern with neutrophils (Figure 3D), while C1, C2, C3 and C5 all have a large part of cells more likely to be iPS cells (Figure 3A, 3B, 3C, 3E).

What’s more, to determine genes which were differential mutated in one subclone, we applied Fisher’s Exact Test on TNBC patients’ mutation details to filter genes which have different mutation frequency between high and low subclone proportions (mean threshold, Table 1). And only differential mutated genes in C4 can be annotated and them enriched in one of the most popular tumor-related pathway - “ErbB signaling pathway” (hsa04012 in KEGG), including ERBB4 and PIK3CA which are key genes in that pathway (Cizkova et al., 2012; Hoque et al., 2010; Lau et al., 2014).

#### C5 can influence patients’ survival outcome

Functional annotation of differential expressed genes showed that C5 was tightly related to “programmed cell death” and “apoptosis” (Figure 2D), and proved the cancer hallmark “resisting cell death”.(Hanahan and Weinberg, 2011) Interestingly, we also found that different stage patients in TCGA TNBC have nearly same level of C5 proportion, and that subclone kept presenting in all stages (Figure 4C). Hence, we assumed that C5 may be the key subclone in TNBC tumors which can make tumor malignant through resisting programmed cell death then induce patients’ poorer survival outcome.

What’s more, C5 also was the only one subclone that its proportion can influence patients’ survival outcome and the TSNE plot of tumor cells distribution showed that in C5 most of cells are malignant. Based on the result of overall survival analysis, threshold is 0.2 (over 0.2 means high risk), and we found that in GSE96058 133 TNBC patients, C5 proportions’ log-rank test *p*-value and HR [95%CI] were 0.0158 and 2.557 [1.160-5.636] (5 years, Figure 5C), 0.040 and 2.557 [1.018-4.612] (10 years, Figure 5D). While in TCGA 80 TNBC patients, we found that there are just only nine patients signed as dead and made it hard to test relationship between C5 proportions and survival outcome. Furthermore, we comprehensively searched for datasets detected by RNA-seq in GEO database with keywords “breast cancer”, “TNBC” or “triple-negative breast cancer”, but we only obtained GSE96058 which is detected by RNA-seq and recorded clinical information (GSE81538 is another SubSeries but do not provide patients’ survival time or survival events).

**Figure 5.**
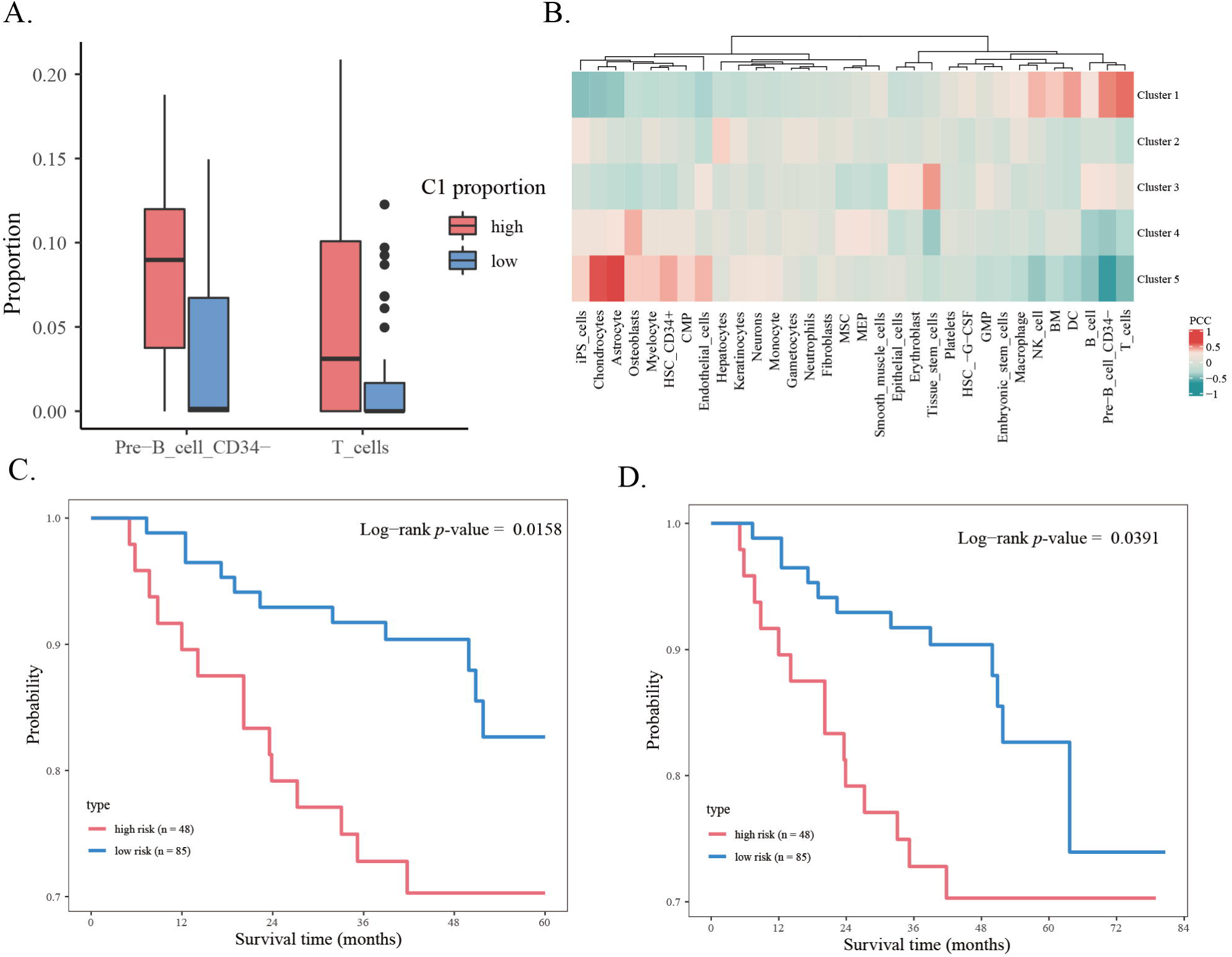
Survival analysis focuses on cluster 5. **A** boxplot for proportions of T cells and pre-B_cell_CD34- in patients with high and low C1 subclonal proportions. **B** heatmap for PCCs among subclones and microenvironment cells. **C-D** K-M plot (5, 10 years overall survival) for cluster 5 proportions in GSE96058 133 TNBC patients and threshold is 0.2.

Taken all together, we found that five subclones all played very important role in tumor development and progression, but on the other hand they also are distinctive with each other. To further verify our hypothesis that: intra-tumor heterogeneity in TNBC embedded in the diversity of the above five subclones, then further brings about inter-tumor heterogeneity reflected in different subclonal composition, we divided TCGA TNBC patients into TNBCtype6 subtypes which have been used wildly (Lehmann et al., 2011). And we found that those six subtypes really have very different subclone composition (Figure 4F), and especially in IM subtype C1 and C2 are enriched markedly which were proofed related to immune functions.

### Microenvironment diversity among TNBC subclones

As analyzed above, we used CIBERSORT algorithm to verify whether immune microenvironments are diverse among TNBC patients, and found that CD4 T cells, Treg cells, Macrophages and mast cells have significant difference among patients with high and low C1 proportions. Although CIBERSORT has become gold standard for immune microenvironment researches, but we hypothesis that if we involve more cell types into analysis then we can estimate more precise cell proportions.

To support our hypothesis, we screened differential expression gene markers of main cell types in HPCA dataset and created simulation data. As described in Materials and Methods part, we got 2823 gene markers for 32 main cell types based on Wilcoxon Rank Sum Test and ssGSVA method (Supplemental table 2). Then we used “RPC” method in “EpiDish” package to test their accuracy and we found that most samples in HPCA have overwhelming proportion of its matched cell type (Supplemental figure 1A). Through comparing cell proportions in these 20 samples between simulation and the standard, we found that comprehensive cell types involved in deconvolution truly can improve the efficiency and accuracy (Supplemental figure 1B). What’s more, we constructed a basis gene expression matrix of the above main cell types which was estimated by mean values.

On the other hand, we believed that more pure microenvironment gene expression profile can avoid the influence of tumor cells. Hence, we proposed a novel method based on cancer purity and genes relative expressing intensities between normal and tumor samples to estimate overall expressing intensities of microenvironment and tumor cells. As described in Materials and Methods part, to estimate the x% which was used to divide samples into purely normal and purely tumor, we set x from 75 to 95 and found that in TCGA BRCA dataset set x as 88 would be the best, in which the purely normal samples have the highest Pearson Correlation Coefficient (PCC) with normal samples and the lowest PCC with tumor samples (Supplemental figure 1B and 1C). Then based on that, we calculated relative gene expressing intensities between tumor and normal and got gene expression profiles for microenvironment and tumor cells through the formula (3) and (4).

Finally, we applied “RPC” deconvolution method on gene expression profiles of microenvironment cells with basic gene expression matrix constructed from HPCA, and we got cell proportions in 83 TNBC samples. Through calculating the PCCs of proportions among subclones and main cell types, we found that C1 is also closely related to T cells and pre-B_cell_CD34- (corresponding PCCs were 0.5240 and 0.4756) which may approve the opinion that C1 is tightly associated with immune-related functions (Figure 5A and 5B).

## Materials and Methods

### Data source

We got the TNBC single-cell RNA-seq data GSE118389 including 1534 cells from GEO database (https://www.ncbi.nlm.nih.gov/geo/) and obtained TCGA data of 113 TNBC patients from GDAC FIREHOSE database (http://gdac.broadinstitute.org) including gene expression, copy number variation, somatic gene mutation data and corresponding clinical information [version: 2016_01_28]. What’s more, to further verify our result, we also downloaded an external dataset GSE96058 which contained RNA-seq data of 135 TNBC patient and corresponding clinical information.

### Cell type classification

Based on the Human Primary Cells Atlas (HPCA) cell expression dataset stored in R package “SingleR” which was named as “hpca.rda”, we used differential expressing gene markers (stored as “hpca$de.genes”) between each two cell type and corresponding expressing data to classify single cells’ cell type (Mabbott et al., 2013). For one unknown single cell and two cell types, we used Euclidean distance to determine that cell is more likely to which cell type; then used Bubbling Sort method to traverse all of the main cell types for identifying the best match cell type for that unknown cell.

### Subclone identification in TNBC

To identify tumor subclones in TNBC cells, we inferred copy number variation (CNV) from RNA-seq gene counts data by using the R package “inferCNV” (https://github.com/broadinstitute/inferCNV) and clustered cells by K-means method. For “inferCNV” part, we used GETx (https://gtexportal.org/home/) samples as the reference normal samples and set default arguments but only sd_amplifier changed as 2. And for cell clustering part, we estimated the best cluster number by “clusGap” function from R package “cluster” with default arguments, then clustered 1000 times randomly (set the seed from 1 to 1000) to definite the final seed which has the lowest tot.withinss and the largest betweenness. Combined with the final seed and the best cluster number, we applied “kmeans” function on cell CNV data with default arguments to distribute every cell to its closest cluster.

### Subclonal gene expression marker identification

After clustering, we got several subclones within TNBC cells which have significant differences on gene copy number. Based on gene TPM values provided by GSE118389, we identified gene expressing markers for each subclone. Firstly, we filtered genes by Analysis of Variance (“anova” function) with threshold of adjusted *p*-value < 0.1 (“p.adjust” function, BH method); then applied Wilcoxon Rank Sum Test (“wilcox.test”, alternative = “greater”) on that highly variable genes, cluster by cluster, to determine cluster-specific gene markers (adjusted *p*-value < 0.05 and Fold Change value > 2). And we used DAVID (https://david-d.ncifcrf.gov/) to annotate subclonal biologic function based on their marker genes. Furthermore, we constructed subclone basis matrix based on the above cluster-specific gene markers and their expression patterns (threshold is mean value).

### Subclonal proportion estimating in bulk tissue

Based on the basis matrix constructed from heterogeneous subclones, we used R package “EpiDISH” to deconvolute bulk gene expressing data (TPM value) to subclonal proportion by CBS deconvolution algorithm. Then we performed the survival analysis on the subclonal proportions, we fitted Cox proportional hazards regression models and *p*-values are obtained from log-rank tests.

### Tissue expression profiles factorization

Based on our hypothesis, we believe that both microenvironment and tumor cells have contributions to tissue expression profiles but their intensities are diversity among genes. Hence, we considered on cancer purity and genes relative expressing intensities between normal and tumor samples to estimate overall expressing intensities of microenvironment and tumor cells.

We collected TCGA breast cancer samples’ purity from the result of Dvir Aran et al. (including purity estimated by ESTIMATE, ABSOLUTE, LUMP, IHC and CPE algorithms) and we used ABSOLUTE as standard because it directly measures tumour cells proportion in a sample (if one sample don’t have ABSOLUTE purity which would be replaced by mean value of the left algorithms). Then we filtered samples of two classes: a) purely normal, including normal samples and patients with low purity lower than x%; b) purely tumor, including patients with high purity more than (1-x)%, and relative genes expressing intensities were defined as Fold Change value of these two classes (mean values of tumor / mean value of normal).

Finally, we can estimate every gene expressing intensities for microenvironment and tumor cells combined with that sample’s tumor purity and genes relative expressing intensities:

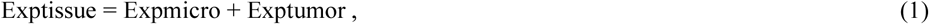

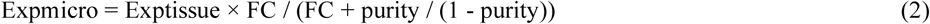

Where Exp means gene expression, FC is Fold Change value of genes between pure normal and pure tumor samples and purity is patients’ tumor purity.

### HPCA cell markers Identification

We collected cell expression dataset from Human Primary Cells Atlas and filtered cell markers of main cell types. For each cell type, we firstly got the union of differential expressing genes (DEGs) from “hpca$de.genes” which only stored DEGs between two cell types; secondly, we applied Wilcoxon Rank Sum Test on those DEGs between this cell class and the others, and only remained genes with p-value < 0.05; finally, we applied ssGSVA on the above DEGs and only kept genes which in samples of that specific cell type have ssGSVA scores all greater than zero. After the above steps, we got marker genes for every main cell types in HPCA dataset.

#### Deconvolution data simulation

To check out whether the number of cell types used in deconvolution would influence the accuracy, we used T cells as the reference to create simulation data. We randomly chose 20 main cell types in HPCA dataset (including T cells) and created 20 samples gene expression profiles with different cell types composition. For these gene expression profiles, we firstly calculated each cell types mean gene expression then based on the cell types proportions to get the final profiles. After that, we randomly chose different number of cell types (must including T cells) to deconvolute these profiles (“RPC” method in EpiDish package), then compared the proportion of T cells in 20 samples between the simulation and the standard.

#### Statistical analysis

In this work we used various statistical tests and algorithms : 1) ssGSVA method was applied by R package “GSVA”; 2) Wilcoxon Rank Sum Test was applied by R function “wilcox.test”; 3) Analysis of Variance (ANOVA) was applied by R function “anova”; and survival analysis was applied by R package “OIsurv”.

## Discussion

Single-cell RNA-seq has offered unprecedented opportunity for deeply understanding the origin, evolution and heterogeneity of cancer, but also brought the challenge in how to associate single-cell genotype with tissue phenotype. Hence, we developed a systematic pipeline for providing a novel insight to understand the subclonal architecture and their biological functions of TNBC through the integration of TNBC scRNA-seq data and deconvolution algorithm. We characterized the inner diversity among TNBC single cells on CNV level and gene expression level then identified five heterogeneous subpopulations. Combined with HPCA reference cell type markers collected in “singleR” package, we identified potentially cell type for every single cell based on gene expression similarity and we found that a great number of cells are more likely to be iPS cells which have highly differentiation pluripotency. After that, based on “inferCNV” method we identified five distinguish cell subpopulations and characterized the biologic functions or their special roles played in cancer progressing. After in-depth and detailed analysis, we found that C1, C2, C4 were related to immune functions, C3 may potential connected with tumor metastasis and C5 might the key subclone in TNBC.

Especially, C1 is tightly associated with the GO term “regulation of T cell activation” and the result of immune microenvironment also shows that C1 can influence the distribution of CD4 T cells, Treg cells, Macrophages and mast cells. And considering on the relationship between C2 and TMB and C4 is enriched in neutrophils, we guess TNBC maybe one special subtype in breast cancer can benefits from immuno-oncology (IO) therapies. On the other hand, our analyze on tumor metastasis shows that not only C3 but also C1, C2 and C5 play important role in that and indicates that metastasis is not the result of single factor but caused by a series of factors. Most importantly, we detect one key subclone C5 which can influence patients’ survival outcome, high C5 proportion means poor outcome. Whatever its enriched GO term “programmed cell death” and its stable distribution in patients with different stage all indicate that C5 may be the hotspot to understand TNBC.

What’s more, we used simulation data proved that comprehensive cell types involved in deconvolution truly can improve the efficiency and accuracy, and if we can get whole cells with different type maybe we can describe tumor environment ecosystem more detailed and penetrating. And we proposed a novel method to split tissue gene expression profiles into microenvironment and tumor which is based on cancer purity and genes relative expressing intensities. Based on the above two parts, we estimated cell proportions in BRCA samples and explored the relationship between microenvironment and subclones, and we found that C1 is closely related to T cells which also approved the enriched GO terms “regulation of T cell activation”.

Taken together, we presented a new thought to analysis tumor heterogeneity and found every subclone plays important but entirely different roles in TNBC and single-cell combined with deconvolution algorithm may be one powerful weapon to understand that deadly cancer type. Additionally, the result of survival analysis applied on C5 let us believe that C5 can be one biomarker for diagnosis and treatment of TNBC. Undoubtedly, future studies will uncover additional features and reveal novel cell subpopulations to understand intra-tumor heterogeneity in tumor progression. Through deconvolution transformed we link single cells and bulk tissue together, and our study substantiates subclonal microenvironment can reveal intra-tumor heterogeneity and influence intra-tumor heterogeneity.

## Figure legends

**Supplemental figure 1.**
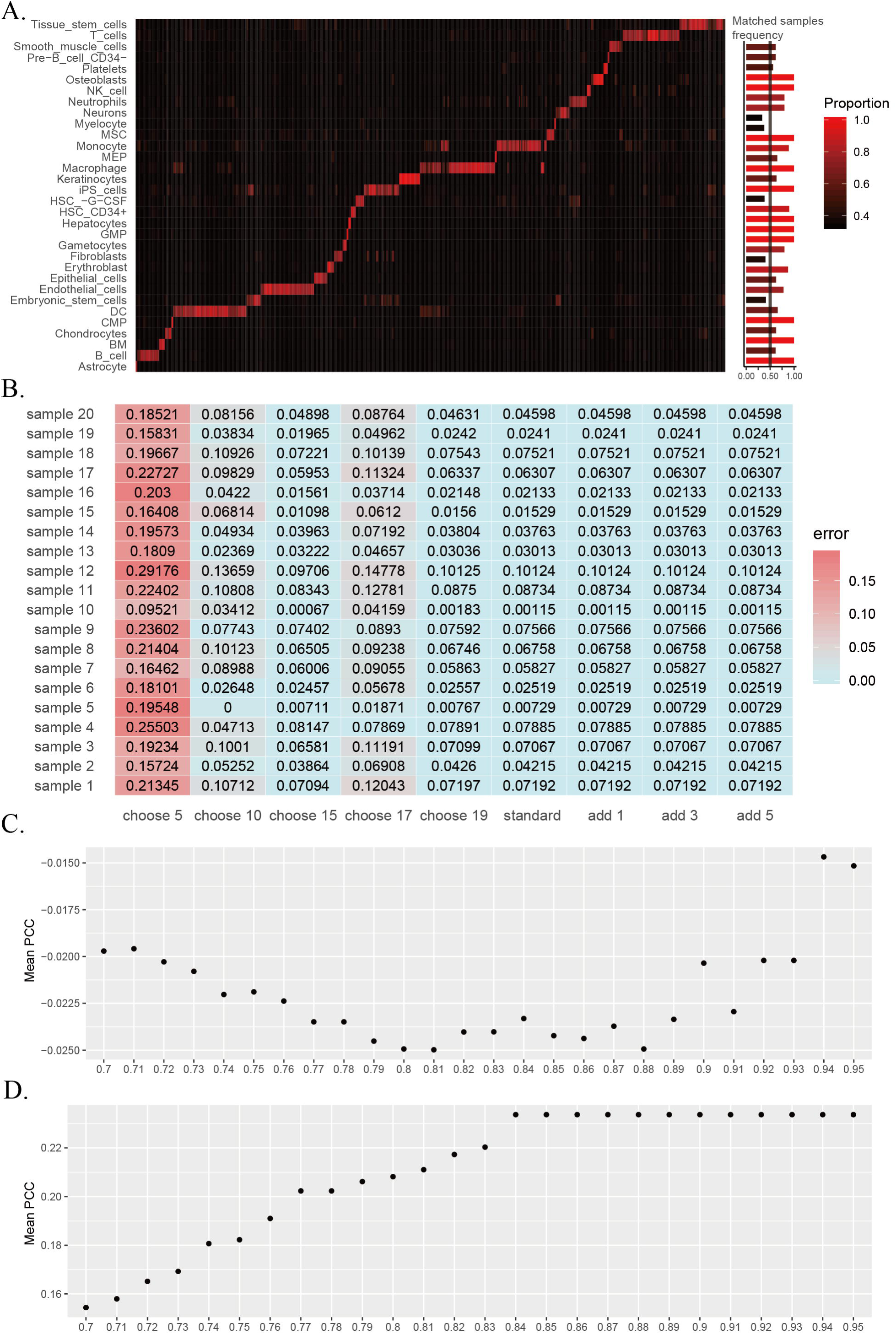
Details for estimating microenvironment. **A** heatmap for HPCA samples’ proportions of main cell types and bar plot for samples of one specific cell type whether that cell’s proportion over 0.6. **B** heatmap for T cells’ proportions among different simulate situations. **C-D** point plots for PCCs to estimate x% used to divide samples to pure normal and pure tumor classes.

## References

Aarts, W.M., Bende, R.J., Bossenbroek, J.G., Pals, S.T., and van Noesel, C.J. (2001). Variable heavy-chain gene analysis of follicular lymphomas: subclone selection rather than clonal evolution over time. Blood 98, 238–240.

Allgauer, M., Budczies, J., Christopoulos, P., Endris, V., Lier, A., Rempel, E., Volckmar, A.L., Kirchner, M., von Winterfeld, M., Leichsenring, J., et al. (2018). Implementing tumor mutational burden (TMB) analysis in routine diagnostics-a primer for molecular pathologists and clinicians. Translational lung cancer research 7, 703–715.

Aran, D., Looney, A.P., Liu, L., Wu, E., Fong, V., Hsu, A., Chak, S., Naikawadi, R.P., Wolters, P.J., Abate, A.R., et al. (2019). Reference-based analysis of lung single-cell sequencing reveals a transitional profibrotic macrophage. Nature immunology 20, 163–172.

Arienti, C., Pignatta, S., Zanoni, M., Cortesi, M., Zamagni, A., Piccinini, F., and Tesei, A. (2017). Looking for Driver Pathways of Acquired Resistance to Targeted Therapy: Drug Resistant Subclone Generation and Sensitivity Restoring by Gene Knock-down. Journal of visualized experiments : JoVE.

Bao, X., Shi, R., Zhang, K., Xin, S., Li, X., Zhao, Y., and Wang, Y. (2019). Immune Landscape of Invasive Ductal Carcinoma Tumor Microenvironment Identifies a Prognostic and Immunotherapeutically Relevant Gene Signature. Frontiers in oncology 9, 903.

Boumber, Y. (2018). Tumor mutational burden (TMB) as a biomarker of response to immunotherapy in small cell lung cancer. Journal of thoracic disease 10, 4689–4693.

Buoncervello, M., Gabriele, L., and Toschi, E. (2019). The Janus Face of Tumor Microenvironment Targeted by Immunotherapy. International journal of molecular sciences 20.

Campa, F., and Randazzo, P.A. (2008). Arf GTPase-activating proteins and their potential role in cell migration and invasion. Cell adhesion & migration 2, 258–262.

Casasent, A.K., Schalck, A., Gao, R., Sei, E., Long, A., Pangburn, W., Casasent, T., Meric-Bernstam, F., Edgerton, M.E., and Navin, N.E. (2018). Multiclonal Invasion in Breast Tumors Identified by Topographic Single Cell Sequencing. Cell 172, 205–217 e212.

Chen, J., Zhou, Q., Wang, Y., and Ning, K. (2016). Single-cell SNP analyses and interpretations based on RNA-Seq data for colon cancer research. Scientific reports 6, 34420.

Cho, E.Y., Chang, M.H., Choi, Y.L., Lee, J.E., Nam, S.J., Yang, J.H., Park, Y.H., Ahn, J.S., and Im, Y.H. (2011). Potential candidate biomarkers for heterogeneity in triple-negative breast cancer (TNBC). Cancer chemotherapy and pharmacology 68, 753–761.

Chu, L.F., Leng, N., Zhang, J., Hou, Z., Mamott, D., Vereide, D.T., Choi, J., Kendziorski, C., Stewart, R., and Thomson, J.A. (2016). Single-cell RNA-seq reveals novel regulators of human embryonic stem cell differentiation to definitive endoderm. Genome biology 17, 173.

Cizkova, M., Susini, A., Vacher, S., Cizeron-Clairac, G., Andrieu, C., Driouch, K., Fourme, E., Lidereau, R., and Bieche, I. (2012). PIK3CA mutation impact on survival in breast cancer patients and in ERalpha, PR and ERBB2-based subgroups. Breast cancer research : BCR 14, R28.

Cresswell, G.D., Apps, J.R., Chagtai, T., Mifsud, B., Bentley, C.C., Maschietto, M., Popov, S.D., Weeks, M.E., Olsen, O.E., Sebire, N.J., et al. (2016). Intra-Tumor Genetic Heterogeneity in Wilms Tumor: Clonal Evolution and Clinical Implications. EBioMedicine 9, 120–129.

Cusnir, M., and Cavalcante, L. (2012). Inter-tumor heterogeneity. Human vaccines & immunotherapeutics 8, 1143–1145.

Ferrone, S., and Whiteside, T.L. (2007). Tumor microenvironment and immune escape. Surgical oncology clinics of North America 16, 755–774, viii.

Garvin, T., Aboukhalil, R., Kendall, J., Baslan, T., Atwal, G.S., Hicks, J., Wigler, M., and Schatz, M.C. (2015). Interactive analysis and assessment of single-cell copy-number variations. Nature methods 12, 1058–1060.

Giessler, K.M., Kleinheinz, K., Huebschmann, D., Balasubramanian, G.P., Dubash, T.D., Dieter, S.M., Siegl, C., Herbst, F., Weber, S., Hoffmann, C.M., et al. (2017). Genetic subclone architecture of tumor clone-initiating cells in colorectal cancer. The Journal of experimental medicine 214, 2073–2088.

Hanahan, D., and Weinberg, R.A. (2011). Hallmarks of cancer: the next generation. Cell 144, 646–674.

Hongu, T., Yamauchi, Y., Funakoshi, Y., Katagiri, N., Ohbayashi, N., and Kanaho, Y. (2016). Pathological functions of the small GTPase Arf6 in cancer progression: Tumor angiogenesis and metastasis. Small GTPases 7, 47–53.

Hoque, M.O., Brait, M., Rosenbaum, E., Poeta, M.L., Pal, P., Begum, S., Dasgupta, S., Carvalho, A.L., Ahrendt, S.A., Westra, W.H., et al. (2010). Genetic and epigenetic analysis of erbB signaling pathway genes in lung cancer. Journal of thoracic oncology : official publication of the International Association for the Study of Lung Cancer 5, 1887–1893.

Hui, T., Cao, Q., Wegrzyn-Woltosz, J., O’Neill, K., Hammond, C.A., Knapp, D., Laks, E., Moksa, M., Aparicio, S., Eaves, C.J., et al. (2018). High-Resolution Single-Cell DNA Methylation Measurements Reveal Epigenetically Distinct Hematopoietic Stem Cell Subpopulations. Stem cell reports 11, 578–592.

Jiang, Y., Xie, J., Huang, W., Chen, H., Xi, S., Han, Z., Huang, L., Lin, T., Zhao, L.Y., Hu, Y.F., et al. (2019). Tumor Immune Microenvironment and Chemosensitivity Signature for Predicting Response to Chemotherapy in Gastric Cancer. Cancer immunology research.

Jin, J., Kong, G.D., and Yoon, H.J. (2018). Deconvolution of Tunneling Current in Large-Area Junctions Formed with Mixed Self-Assembled Monolayers. The journal of physical chemistry letters 9, 4578–4583.

Karaayvaz, M., Cristea, S., Gillespie, S.M., Patel, A.P., Mylvaganam, R., Luo, C.C., Specht, M.C., Bernstein, B.E., Michor, F., and Ellisen, L.W. (2018). Unravelling subclonal heterogeneity and aggressive disease states in TNBC through single-cell RNA-seq. Nature communications 9, 3588.

Lachapelle, J., and Foulkes, W.D. (2011). Triple-negative and basal-like breast cancer: implications for oncologists. Current oncology 18, 161–164.

Lau, C., Killian, K.J., Samuels, Y., and Rudloff, U. (2014). ERBB4 mutation analysis: emerging molecular target for melanoma treatment. Methods in molecular biology 1102, 461–480.

Lehmann, B.D., Bauer, J.A., Chen, X., Sanders, M.E., Chakravarthy, A.B., Shyr, Y., and Pietenpol, J.A. (2011). Identification of human triple-negative breast cancer subtypes and preclinical models for selection of targeted therapies. The Journal of clinical investigation 121, 2750–2767.

Li, Z., Ren, M., Tian, J., Jiang, S., Liu, Y., Zhang, L., Wang, Z., Song, Q., Liu, C., and Wu, T. (2015). The differences in ultrasound and clinicopathological features between basal-like and normal-like subtypes of triple negative breast cancer. PloS one 10, e0114820.

Liu, Z., Li, M., Jiang, Z., and Wang, X. (2018). A Comprehensive Immunologic Portrait of Triple-Negative Breast Cancer. Translational oncology 11, 311–329.

Mabbott, N.A., Baillie, J.K., Brown, H., Freeman, T.C., and Hume, D.A. (2013). An expression atlas of human primary cells: inference of gene function from coexpression networks. BMC genomics 14, 632.

Malchenko, S., Galat, V., Seftor, E.A., Vanin, E.F., Costa, F.F., Seftor, R.E., Soares, M.B., and Hendrix, M.J. (2010). Cancer hallmarks in induced pluripotent cells: new insights. Journal of cellular physiology 225, 390–393.

Marra, A., Viale, G., and Curigliano, G. (2019). Recent advances in triple negative breast cancer: the immunotherapy era. BMC medicine 17, 90.

Mills, M.N., Yang, G.Q., Oliver, D.E., Liveringhouse, C.L., Ahmed, K.A., Orman, A.G., Laronga, C., Hoover, S.J., Khakpour, N., Costa, R.L.B., et al. (2018). Histologic heterogeneity of triple negative breast cancer: A National Cancer Centre Database analysis. European journal of cancer 98, 48–58.

Montagna, E., Maisonneuve, P., Rotmensz, N., Cancello, G., Iorfida, M., Balduzzi, A., Galimberti, V., Veronesi, P., Luini, A., Pruneri, G., et al. (2013). Heterogeneity of triple-negative breast cancer: histologic subtyping to inform the outcome. Clinical breast cancer 13, 31–39.

Nault, J.C. (2014). Next generation sequencing, inter-tumor heterogeneity and prognosis of hepatitis B related hepatocellular carcinoma. Chinese journal of cancer research = Chung-kuo yen cheng yen chiu 26, 730–731.

Newcomb, E.W., Silverstein, S.C., and Silagi, S. (1978). Malignant mouse melanoma cells do not form tumors when mixed with cells of a non-malignant subclone: relationships between plasminogen activator expression by the tumor cells and the host’s immune response. Journal of cellular physiology 95, 169–177.

Newman, A.M., Liu, C.L., Green, M.R., Gentles, A.J., Feng, W., Xu, Y., Hoang, C.D., Diehn, M., and Alizadeh, A.A. (2015). Robust enumeration of cell subsets from tissue expression profiles. Nature methods 12, 453–457.

Poirion, O., Zhu, X., Ching, T., and Garmire, L.X. (2018). Using single nucleotide variations in single-cell RNA-seq to identify subpopulations and genotype-phenotype linkage. Nature communications 9, 4892.

Prat, A., Adamo, B., Cheang, M.C., Anders, C.K., Carey, L.A., and Perou, C.M. (2013). Molecular characterization of basal-like and non-basal-like triple-negative breast cancer. The oncologist 18, 123–133.

Romero, D. (2019). TMB is linked with prognosis. Nature reviews Clinical oncology 16, 336.

Schelker, M., Feau, S., Du, J., Ranu, N., Klipp, E., MacBeath, G., Schoeberl, B., and Raue, A. (2017). Estimation of immune cell content in tumour tissue using single-cell RNA-seq data. Nature communications 8, 2032.

Stanta, G., and Bonin, S. (2018). Overview on Clinical Relevance of Intra-Tumor Heterogeneity. Frontiers in medicine 5, 85.

Werbowetski-Ogilvie, T.E., and Bhatia, M. (2008). Pluripotent human stem cell lines: what we can learn about cancer initiation. Trends in molecular medicine 14, 323–332.

Wright, N., Rida, P.C.G., and Aneja, R. (2017). Tackling intra- and inter-tumor heterogeneity to combat triple negative breast cancer. Frontiers in bioscience 22, 1549–1580.

Wu, Y., Zheng, J., Li, Z., Zhao, Y., and Zhang, Y. (2009). A novel reagentless amperometric immunosensor based on gold nanoparticles/TMB/Nafion-modified electrode. Biosensors & bioelectronics 24, 1389–1393.

Zeng, W., Chen, X., Duren, Z., Wang, Y., Jiang, R., and Wong, W.H. (2019). DC3 is a method for deconvolution and coupled clustering from bulk and single-cell genomics data. Nature communications 10, 4613.

Zheng, B., Wang, J., Cai, W., Lao, I., Shi, Y., Luo, X., and Yan, W. (2019). Changes in the tumor immune microenvironment in resected recurrent soft tissue sarcomas. Annals of translational medicine 7, 387.

Zheng, J., and Gao, P. (2019). Toward Normalization of the Tumor Microenvironment for Cancer Therapy. Integrative cancer therapies 18, 1534735419862352.

Zong, C.C. (2017). Single-cell RNA-seq study determines the ontogeny of macrophages in glioblastomas. Genome biology 18, 235.

