## Supplementary material for "Single-cell RNA-seq data reveals TNBC tumor heterogeneity through characterizing subclone compositions and proportions": Table 1

**Table 1**. Differential mutated genes among five subclones

| **subclone** | **genes** |
| --- | --- |
| Cluster 1 | CARD11; FAM75D5 |
| Cluster 2 | ANK2; BCORL1; CACNA1E; COL18A1; CYP26B1; DLC1; FAM65C; GCC2; HUWE1; KIF24; MUC17; NBAS; NEB; PREX1; RELN; RYR3; TCHH; USP34; UTP20; ZFHX4 |
| Cluster 3 | CSMD3; GBF1 |
| Cluster 4 | CKAP5; ERBB4; FAM86B2; FAM98A; FRMD7; PIK3CA; SCN1A; SUGP1 |
| Cluster 5 | DNAH14; FCGBP; MUC16; SDHAP2; SRRM2; ZNF318; ZNF687 |
