## Supplementary material for "Single-cell RNA-seq data reveals TNBC tumor heterogeneity through characterizing subclone compositions and proportions": Table 2

**Table 2**. Patients details for Cumulative Hypergeometric Test.

|  | low risk patients | high risk patients | All patients |
| --- | --- | --- | --- |
| Non-Metastatic patients | 48 | 10 | 58 |
| Metastatic patients | 16 | 9 | 25 |
| All patients | 64 | 19 | 83 |

\# In R: phyper((48-1), 58, 25, 64, lower.tail = F) = 0.059.
